# Repeated blood-brain barrier opening using low-intensity pulsed ultrasound mitigates amyloid pathology

**DOI:** 10.1101/2025.04.28.650998

**Authors:** Michael Canney, Guillaume Bouchoux, Alexandre Carpentier, Pierre De Rossi

## Abstract

**Background:** The delivery of large molecules to the pathological brain is one of the main obstacles in the development of disease-modifying drugs. This is partly due to the presence of the blood-brain barrier (BBB), which blocks the free passage of lipophobic molecules and those larger than 400 Da. One strategy to bypass this natural barrier is to use low intensity pulsed ultrasound (LIPU) to oscillate circulating micro-sized microbubbles which then exert mechanical stress on the vessel walls. This procedure allows for temporary disruption of the BBB and enhanced local delivery of therapeutics from the blood to the brain parenchyma. In this study, the effect of repeated BBB opening on neuroinflammation in a healthy mouse model was first explored followed by the effect of repeated opening on amyloid pathology in a model for Alzheimer’s disease.

**Methods:** A cohort of wildtype mice was used to determine the effect of a single BBB opening session mediated by ultrasound/microbubbles (US/MB) on the inflammatory profile, using RT-Q-PCR on brain extracts at 2-, 4-, 8- and 15-days post opening. A second cohort of ARTE10 mice, a mouse model for amyloid pathology, was treated with a different sequence of repeated US/MB mediated BBB opening to explore the effect on the pathology. Tissues were also analyzed for immune cell infiltration, microglia and astrocyte activation, and inflammatory response.

**Results:** Our results demonstrate that the opening of the BBB leads to a mild inflammatory response in wild-type animals. However, repeated opening of the BBB in the ARTE10 model resulted in a mild decrease in amyloid pathology, along with a mild increase of growth factor.

**Conclusions:** Altogether, this study suggests that sonication is not only a safe method to deliver therapeutics to the brain but could also have synergistic effects in the treatment of neurodegenerative diseases.

## Introduction

Alzheimer’s disease (AD) is a neurodegenerative disorder that slowly affects memory, thinking and behavior, associated with progressive neurodegeneration. AD is characterized by two molecular hallmarks: the extracellular aggregation of amyloid-beta in plaques and the intracellular aggregation of hyperphosphorylated tau, resulting in tau tangles. These progressive pathological features have been hypothesized to lead to the resulting cognitive deterioration observed in patients.

While the last two decades of research have uncovered several risk factors for AD and led to the discovery of new pathways that may be responsible for disease onset, most pharmaceutical development has been focused on developing therapies targeting tau and amyloid. The hypothesis behind this strategy is that the removal of these pathological aggregates will lead to restoration of normal neuronal function and delay or stop the progression of the disease.

Recently, the first regulatory approvals have been obtained for antibody-based therapies targeting amyloid for the treatment of AD, despite only showing modest results in Phase 3 clinical trials[1]. However, one of the main obstacles that limit the efficacy of these antibody treatments targeting amyloid is the difficulty in bypassing the blood-brain barrier (BBB). Normally, only 0.01%[2] of the administered antibody reaches the brain. If higher parenchymal concentrations could be reached, we could observe a better clearance of the plaques associated with a better efficacy of these treatments, as recently demonstrated by the last results for the phase 2 clinical trial for the Trontinemab[3].

To overcome the BBB, ultrasound (US) technology can be used to temporarily disrupt the BBB and increase delivery of therapeutics. This method uses the combination of circulating/oscillating microbubbles (MB) administered intravenously and stimulated by low intensity pulsed ultrasound (LIPU) to disrupt the tight junctions of the BBB, allowing enhanced entry of molecules, ranging from small molecule therapeutics to large antibodies into the parenchyma[4–21]. This strategy has been used to enhance delivery of therapeutics into the CNS of mouse models for AD[4,7,9]. In addition, a recent clinical trial by Rezai et al. 2024[15] has shown promising results for delivering the FDA-approved anti-amyloid therapy aducanumab in Alzheimer’s patients using US/MB with a clinical ultrasound system. A 32% reduction in SUVR on ^18^F-florbetaben PET scans was observed after 26 weeks in the regions that received treatment to open the BBB. These clinical results demonstrate the potential efficacy gained by the delivery of antibody therapies using ultrasound.

US-mediated BBB opening alone, without concomitant drug administration has also been shown to have beneficial effects, potentially by altering the immune microenvironment in Alzheimer’s mouse models and in patients[4,22–30], with improved outcomes when coupled to anti-amyloid therapies[4,31–35]. Moreover, it was shown that opening of the BBB can also potentiate a diverse range of beneficial biological effects including neurogenesis[36–38]. In amyloid models of the disease, repeated BBB opening decreased the number of amyloid deposits in the brain[22,24,27–30,35] and improved cognition[22,24,27,30]. Similarly, in tau models of the disease, repeated BBB opening was able to decrease tau pathology[39,40] and improve cognition[39]. Finally, in patients with AD, repeated opening of the BBB using US was shown to be safe[41–44] and efficient to mitigate the amyloid burden in the brain[43,44].

While several studies have already explored the effect of BBB opening using US/MB on the inflammatory profile of the brain at short term, evidence is still lacking on the lasting effects of BBB opening in the healthy brain as well as its potential as a therapeutic approach for neurodegenerative diseases. In this pilot study, we explored the effect of BBB opening in wildtype mice and in a mouse model for amyloid pathology (ARTE10) on the inflammatory profile of the brain and its effect on neuropathological hallmarks.

## Experimental Procedures

### Mouse models

Twenty (20) male C57BL/6J mice of approximately 17 weeks of age were obtained from The Jackson Laboratory (Bar Harbor, ME). Thirty-six (36) one-year-old females of the ARTE10 mouse model were used as a model for amyloid pathology and obtained from Taconic.

### Sonication

The following ultrasound parameters were used: 120Lseconds using a 1-MHz 25,000-cycle burst at a 1LHz pulse repetition frequency and an acoustic pressure of 0.3LMPa (Carthera, Lyon, France). To prepare for sonication, the hair on the head of all animals was removed using depilatory cream. At the time of sonication, mice were injected intraperitoneally with 5 μL/gram of 90 mg/kg ketamine / 10 mg/kg xylazine (Zetamine: VetOne, catalogue No. AH202C6)/xylazine (Rompun: Bayer, catalogue No. AH200W5). Lumason microbubbles (Bracco Diagnostics, catalogue No. 20A108A) were injected into the retro-orbital sinus (r.o.) at a dose of 150 μL/mouse and mice underwent a single sonication event. Mice were given 10 mg/kg atipamezole (Rivertidine; Modern Veterinary Therapeutics, Cat #023575) i.p. within 20 min of ketamine administration to reverse the anesthesia. After 15 min, blood was collected for PK.

### Experimental Design for WT animals

Animals were weighed on study days 0, 1, 7, and 14. A pre-sonication blood sample was collected on study day 0, as a baseline reference. On study day 1, the animals received a single sonication event. All animals were bled by retro-orbital blood drawn for plasma (EDTA) on study day 0 (baseline; 120 μL of plasma/mouse). Plasma samples were stored frozen at −80°C for analysis of S100. The animals were euthanized for necropsy and terminal sample collection 1 day, 3 days, 7 days, or 14 days post-sonication.

### Experimental Design for ARTE10 animals

Animals were weighed and clinical observations were recorded on D0, D7, D14, D21, D28, and D49. A pre-sonication blood sample was collected on study day 0. Groups 1 and 2 did not receive any sonication (control groups). Groups 3 and 5 (sparse sonication groups, 4 sonication total) received repeated sonication every 2 weeks for 8 weeks with respective sacrifice 1 week and 1 month after the last sonication procedure. Group 4 (dense sonication group, 8 sonication total) received repeated sonication twice a week for 4 weeks and were sacrificed 1 week after the last sonication. Group 6 (very sparse sonication group, 2 sonication total) received repeated sonication every 4 weeks for 8 weeks and were sacrificed 1 week after the last sonication.

### Postmortem brain tissue collection

Brains were halved along the longitudinal fissure. Right brain hemispheres were fixed in 4% PFA, then transferred to 40% sucrose/saline the next day and stored at 4°C for histology. Left brain hemispheres were collected into RNase/DNase free clean tubes and stored at −80°C for gene expression analysis. Cerebrospinal fluid (CSF) samples were collected and whole blood was collected and processed to plasma (EDTA, 300 μL/mouse). Plasma and CSF samples (5-10 μL/mouse) were stored at −80°C for analysis.

### S100-**β** ELISA Methods

Plasma samples were analyzed for S100-β using a Mouse Protein S100-β /S100 Beta ELISA Kit (CUSABio, catalogue No. CSB-EL020643MO, lot No. J16147391). All reagents, working standards, and samples were prepared as directed in the kit instructions.

### Real-Time Quantitative Polymerase Chain Reaction (RT-qPCR) Methods

#### For WT animals

RT-qPCR analysis was performed on left brain hemispheres from all animals. Tissue samples (approx. 30 mg each) were weighed out and homogenized in RLT buffer on a FastPrep-24 5G. RNA was isolated using a Qiagen RNeasy Mini kit (No. 74106) and the manufacturer-recommended protocol, and RNA was reverse transcribed into cDNA using the ThermoFisher High-Capacity cDNA kit (ThermoFisher, No. 4368814). cDNA samples were then analyzed using Taqman primers against *cyc1*, *cxcl1*, *ccl2*, *Il-1β*, *Il-6*, *tnf-a*, *Icam*, *Il-10*, *tgf-β*, *Il-13*, *bdnf*, *nf-κb*, *p-Selectin*, *e-Selectin*, GM-CSF, and G-CSF. All primers were selected from validated primer sets from ThermoFisher Scientific.

#### For ARTE10 cohort

Multiplex qPCR master mix was prepared using TaqMan Multiplex Master Mix (ThermoFisher, Cat #4484263) according to manufacturer’s specifications for 96-well plates. Samples were run in triplicate for the following genes: *Il-1*β*, Il-6, Il-10, tnf-*α*, bdnf, ccl2, Icam1*

#### Histological Processing

Mouse hemibrains were embedded in paraffin, and brain sections were mounted on glass slides and stained with hematoxylin and eosin (H&E) using standard methods. For immunohistochemical (IHC) staining of mouse hemibrains, staining was conducted on the Leica Bond RXm platform using standard chromogenic methods. For antibodies requiring heat induced epitope retrieval (HIER), slides were heated in a pH9 ethylenediaminetetraacetic acid (EDTA)-based buffer at 94 °C for 25 min.

### Antibody Information

Antibody Information.
| Target | Host | Clone | Vendor | Cat # |
| --- | --- | --- | --- | --- |
| Iba1 | Rabbit | EPR16588 | AbCam | ab178846PUR |
| GFAP | Rabbit | EPR1034Y | AbCam | ab68428 |
| Aβ | Rabbit | D12B2 | Cell Signaling | 9888 |
| CD3 | Rabbit | SP7 | AbCam | ab16669 |

### Staining Parameters

Staining Parameters.
| Antibody | Antigen Retrieval | Dilution | Chromogen |
| --- | --- | --- | --- |
| Iba1 | pH9 | 1:40,000 | DAB |
| GFAP | pH9 | 1:200 | DAB |
| Aβ | pH9 | 1:200 | DAB |
| CD3 | pH9 | 1:100 | DAB |

### Aβ Image Analysis

IHC-stained glass slides (n = 62) labelled for Aβ were scanned using an Aperio AT2 whole slide scanner with a maximum objective of 20x. Image analysis was conducted on Aβ IHC-stained digital slide images using Visiopharm® software. Six coronal sections of the brain per animal (across 2 slides) were annotated manually as individual regions of interest (ROIs): rostral 1 and 2, middle 1 and 2, and caudal 1 and 2. IHC-positive area and total area within each ROI were used to calculate percent positive area for Aβ immunolabeling. The calculations for percentage positive area (%) are provided below:

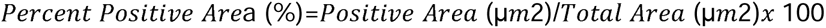

For further analysis a deep learning algorithm trained to detect tissue was run on each slide to define the region of interest (ROI). Deep learning algorithms (DLA) use convolutional neural networks (CNNs) that can be trained to identify specific objects within the tissue based on morphological patterns, RGB signatures, texture, size, and spatial relationships. This algorithm was developed using the U-Net model. A second DLA trained to detect vascular endothelium trained on slides stained with β-amyloid IHC was used to define vessels. These vessels were then outlined into a separate ROI. β-amyloid immunolabeling was detected using a final DLA. β-amyloid signal was assessed as total, plaque and non-plaque. Total β-amyloid burden was defined by the sum of the areas of plaques and non-plaques. For the frontal cortex segmentation, data were reported as percent area of total β-amyloid, percent area of plaques, number of plaques, and average plaque size. These data were reported from the total brain area, including all 6 sections of the brain including vasculature, as well as frontal cortex as defined by the Allen Brain Atlas. Three of the 6 sections had frontal cortex present.

### CD3, GFAP, and IBA1 Image Analysis

CD3, GFAP, and IBA1 immunolabeling were detected using thresholding techniques. All three thresholds were developed individually based on Visiopharm’s proprietary HDAB filter, which isolates the specific intensity and colorimetric signature associated with positive DAB-based immunolabeling. Total brain area was defined as the sum of the non-frontal cortex brain area and the frontal cortex. CD3 quantification is reported as positive cell density/mm2; GFAP and IBA1 quantification are reported as positive percentage area.

### Statistical Analyses

Statistical analyses were performed on the data generated from this study using GraphPad Prism 10.03 for Windows (GraphPad Software, Inc.). Details of each analysis can be found in the figure legend presenting the data. Statistical significance was set at p < 0.05. Considering the sample size of our groups, data showing a statistical trend are also presented in this study.

## Results

### Unique BBB disruption changes the molecular inflammatory signaling profile for up to 15 days post-sonication

To better understand the effects of mechanical opening of the BBB by US/MB, we investigated blood-based markers and inflammatory markers following a single session of ultrasound to disrupt the BBB (experimental plan, Fig 1A).

**Figure 1:**
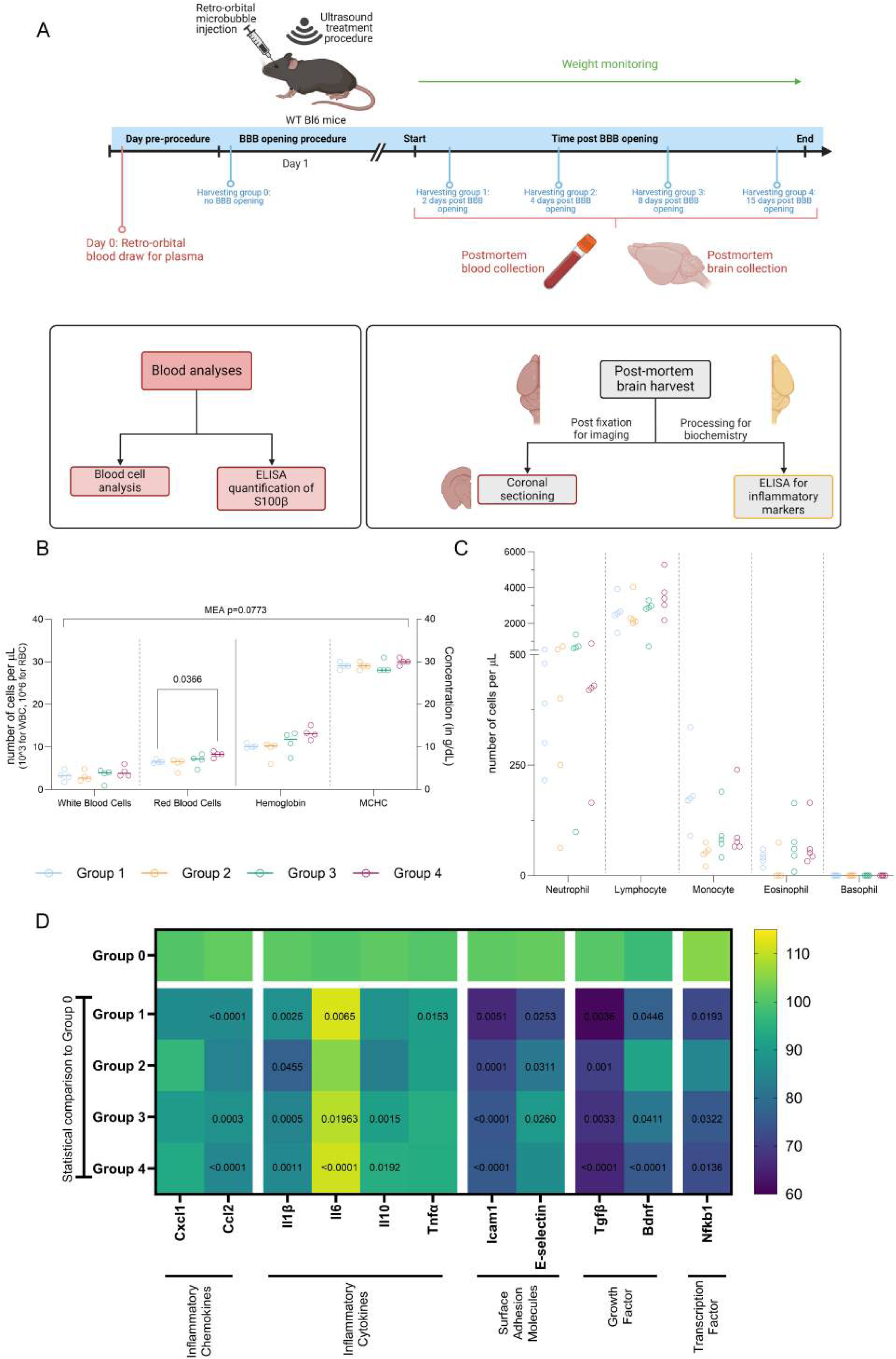
Physiological impact of a single session of ultrasound mediated BBB-opening: **A)** Schematic summarizing the key steps of the protocol. Briefly, wildtype mice were first weighed and a blood sample was collected before undergoing a single sonication procedure. Four groups of mice were followed with terminal points at 2-, 4-, 8- and 15-days post sonication. During this time, mouse weights were monitored. At sacrifice, a blood sample was collected, and brains were harvested for *ex vivo* analyses. **B)** Graphic plots representing the complete blood count, each dot representing one animal. Red blood and White blood cells number were counted in the final collected blood sample (number of cells expressed in nx10^3 for white blood cells and nx10^6 for red blood cells, left Y axis). Hemoglobin levels and Mean corpuscular hemoglobin concentration (MCHC) was measured (in ng/dl, right Y axis). 2-way ANOVA with post hoc Tukey’s multiple comparison test, difference between groups F (3, 12) = 2.923, p=0.0773 **C)** Graphic plots representing the complete blood count, each dot representing one animal. Number of neutrophils, lymphocytes, monocytes, eosinophils and basophils were counted for each group (in cells per microliter). 2-way ANOVA with post hoc Tukey’s multiple comparison test, difference between groups F (12, 48) = 0.8972, P=0.5560 **D)** Heatmap representing the RT-Q-PCR quantification performed on extracted brains for different markers and normalized by the baseline group (group 0). The legend on the left indicates the level of increase or decrease compared to the Mean of the group 0. Colors are based on the Median. P value in each cell indicates the significance compared to the baseline group 0, mixed effect analysis with post hoc dunnett’s multiple comparison test, difference between groups F (4, 18) = 25.68, p<0.0001.

No significant changes in body weight were observed, however with a trend toward weight loss in the first 24h post treatment (ANOVA p=0.06277, D0 vs D1 p=0.0023) (Fig S1A). Treated mice showed a trend towards an increased in red blood cell count 15 days post opening (MEA p=0.0773, group1 vs Group4 p=0.0366) (Fig 1B), however no significant effect was recorded on the number of circulating white cells (Fig 1C). We used the S100b marker to monitor the opening of the BBB post procedure and its level overtime. As expected, we observed a drop of the S100b level over time reaching the lowest level 8 days after BBB opening (LS mean = 16.00), becoming significant only at 15d post sonication (LS mean 16.41) (MEA p=0.0063, post sonication vs terminal 15 days p= 0.0115) (Fig S1B, measured by ELISA).

Once the safety of BBB opening was established, we performed RT-Q-PCR to explore the level of different inflammatory markers in the brain at several time points [2-, 4-, 8- and 15-days post opening] post BBB opening. Interestingly, the only significant increase compared to baseline, group 0 without BBB opening, was an increase in *Il-6* mRNA levels, (MEA p<0.0001, Group 0 vs 1: p=0.0065, Group 0 vs 3 p=0.0003, Group 0 vs 4 p<0.0001). It is important to note that *Il-6* increase was correlated with an increase of red blood cells (Fig 1B)[45]. Notably, all the other factors measured were decreased, either transiently, or with a prolonged effect, up to 15 days post treatment.

Inflammatory chemokines *Cxcl1* and *Ccl2* as well as inflammatory cytokines *Il-1b*, *Il-10* and *Tnf-a* were decreased. While most of the chemokines decreased over 15 days post treatment, *Tnf-a* was only decreased in the first 48h of the protocol, with a recovery to baseline by day 4. Adhesion molecules *Icam1* and *E-selectin* were also decreased, with a recovery of *E-selectin* by day 15. The growth factors *Tgfb* and *Bdnf* were found to be significantly reduced after a single sonication. Finally, the transcription factor *Nfkb* was significantly decreased post sonication. Altogether, these results suggested that the opening of the BBB with the selected parameters did not lead to a pro-inflammatory response, or any significant changes to other physiological metrics in a healthy mouse model.

### Different schedules of repeated BBB opening did not impact physiological output

Next, the effect of different sonication protocols was explored in a mouse model for AD that develops amyloid deposition (ARTE10). We tested different regimens of US/MB mediated BBB opening going from twice a week to once a month with sacrifice and profiling of different markers at one week or one month after the last sonication (Fig.2A). We first investigated the effect of repeated sonication on the general health and survival of the mice. Our data showed no statistical difference between the groups (MEA, p=0.6513), with however a significant difference between timepoints (MEA, p=0.0459, D21 vs D28 p=0.0280), as it was described for the WT cohort. (Fig. 2B). A few mice had weight reduction following the first session of the sonication but recovered in the following week(s).

**Figure 2:**
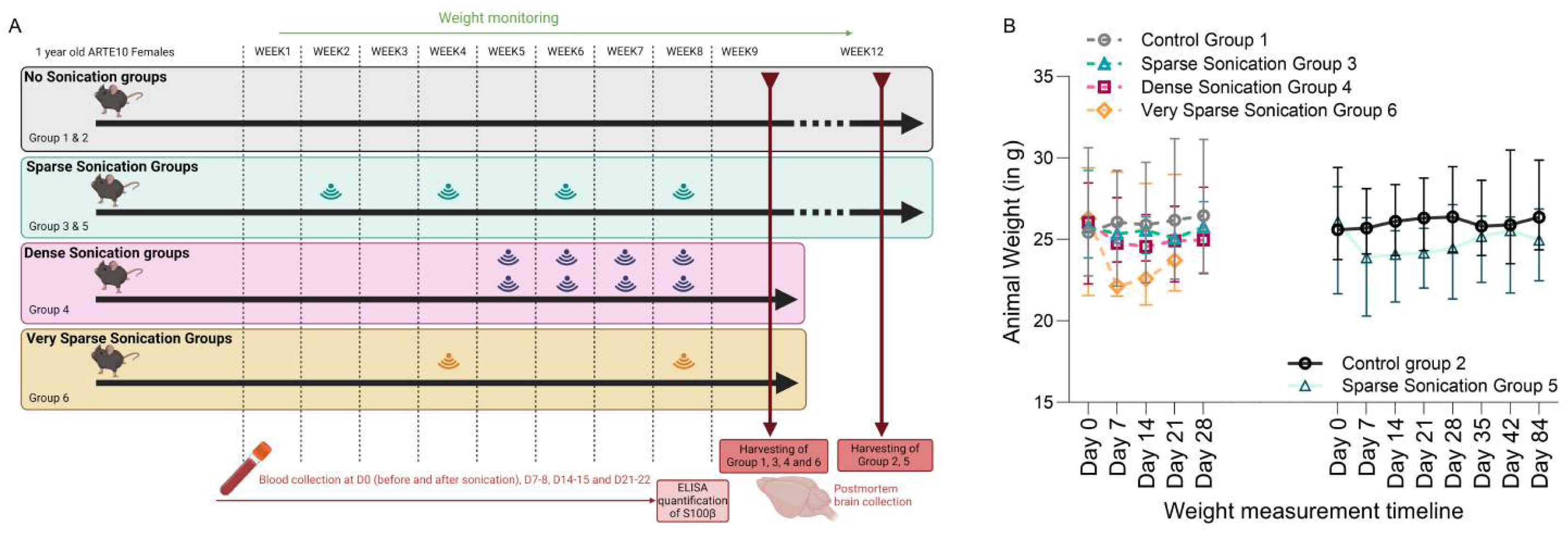
Physiological data on the ARTE10 cohort following repeated sonication protocols: **A)** Schematic summarizing the experimental plan for the different groups (1 to 6), the sonication schedule and sacrificing date following the last sonication session. **B)** Line graph representing the evolution of the median weight for each group (bars represent the range). No significance was observed between groups, mixed effects analysis at 1 week: F (3, 20) = 0.5542, p=0.6513, at 1 month: F (1, 10) = 3.214, p=0.1033.

Overall, our data suggests that LIPU-mediated BBB opening is a safe procedure that allows repetitive treatment. Altogether, these data demonstrated the low level of toxicity of a repeated protocol of sonication, as well as the structural plasticity of the BBB allowing repeated mechanical disruption of its structure.

### Sonication frequency does not trigger inflammation in the brain over time

Using the post-mortem tissues from the ARTE10 cohort, we then investigated the effect of repeated BBB opening on inflammatory markers of the brain at the cellular and the molecular level. CD3, GFAP and Iba1 markers were used to explore T-cell infiltration, gliosis and microglial activation respectively (Fig. 3A). We used an automated pipeline of analysis, able to recognize the cells/markers of interest to extract the number of cells per mm^2^ or the percentage of occupancy (see Supp Fig2. A-C showing the quantification mask for the 3 markers). The obtained results showed that repeated sonication did not lead to an increase of CD3+ cell infiltration (Fig3B and C). A statistical trend toward an increase in astrogliosis was observed between Group 1 and Group 3 (ANOVA p<0.0001, Group 1 vs Group 3 P=0.0918) (Fig3D and F), as previously reported[46] and consistent with the level of S100b observed. In all the other conditions, as well as for all the conditions for Iba1, no difference between groups was detected (Fig3D-F). A few CD3+ cells were observed in the vicinity of amyloid deposits. As expected, GFAP+ and Iba+ cells tend to accumulate around the amyloid deposits. However, to explore how sonication stimulates the relocalization of inflammatory cells around the pathological aggregates, to possibly participate in their elimination, more in-depth studies will be required, to explore the cellular mechanisms triggered by LIPU-mediated BBB opening.

**Figure 3:**
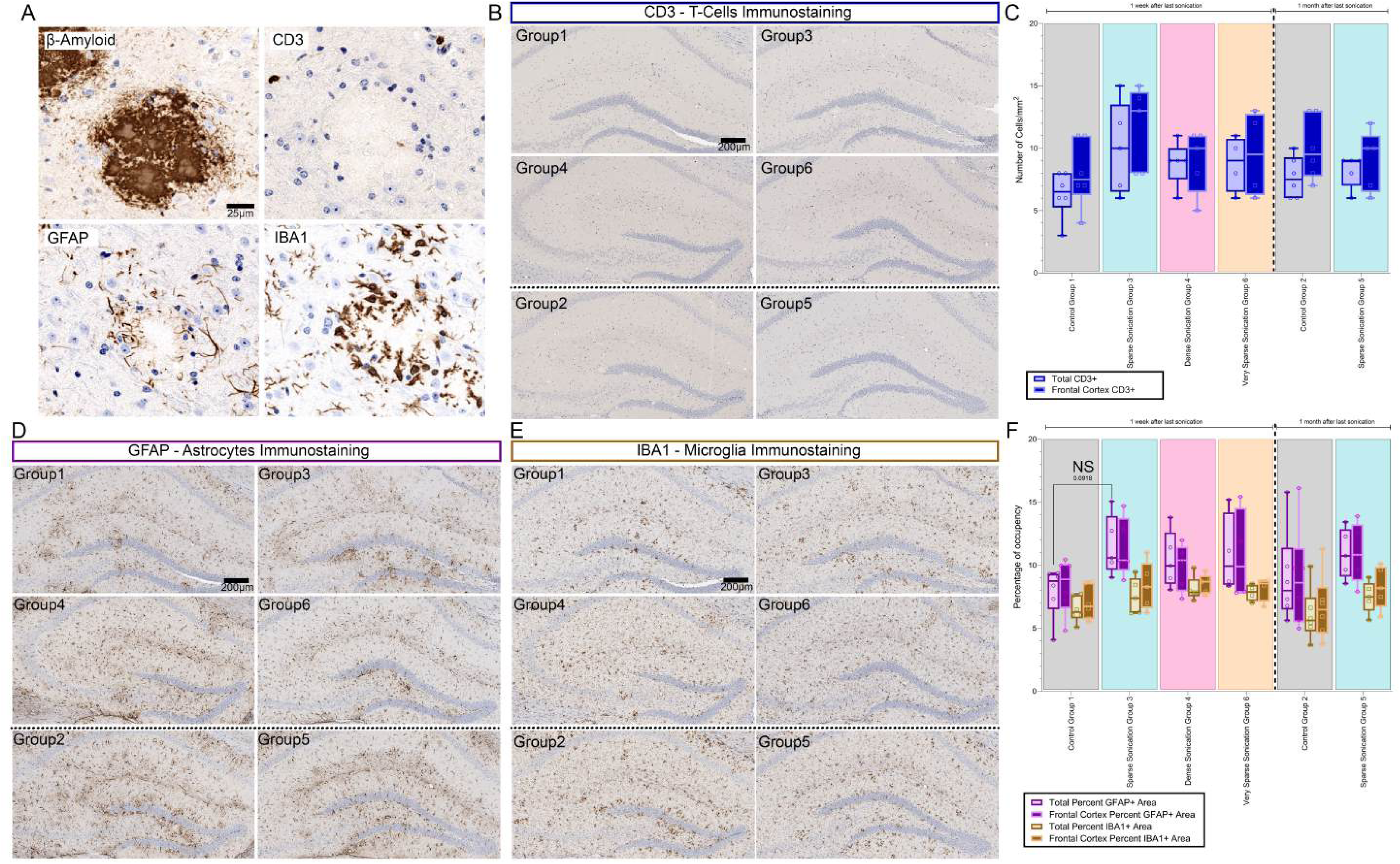
Inflammatory cellular landscape of the brain following repeated sonication procedures: **A)** Representative image showing the presence of CD3+, GFAP+ and Iba1+ cells around an amyloid deposit, labelled with a b-amyloid antibody. **B)** Representative images showing the presence of T-cells (CD3+) present in the brain parenchyma of the different groups. **C)** Box plots showing median, 25 and 75 percentiles and min/max values, along with individual data point for each animal, for the density of T-cells (CD3+) present in the brain parenchyma of the different group. No significant differences between groups were recorded, 2-way ANOVA with post hoc Tukey’s multiple comparisons test, difference between groups F (5, 25) = 1.285, P=0.3016. **D)** Representative images showing the occupancy of astrocytes (GFAP+) in the brain parenchyma of the different groups. **E)** Representative images showing the occupancy of microglial (Iba1+) in the brain parenchyma of the different groups. **F)** Box plot showing median, 25 and 75 percentiles and min/max values, along with individual data point for each animal, for the total occupancy for astrocytes (purple bars) and microglia (yellow bars) in the brain parenchyma of the different groups. No significant differences between groups were recorded, 2-way ANOVA with post hoc Tukey’s multiple comparisons test, difference between groups F (5, 25) = 1.280, P=0.3038.

We generated pilot results on the mRNA expression of few inflammatory markers (as explored in Fig. 1). Interestingly, our data showed a decrease in the proinflammatory cytokines in all conditions, as was observed in wildtype mice (Fig. 1). Opposite to what was observed in the single BBB opening protocol, repeated opening of the BBB was associated with an increase of *bdnf* in most treatments, however not statistically tested due to limited number of data (Supp fig 2D). This brings the possibility that the repeated mechanical opening of the BBB could favor the local release of growth factors, as it was observed in other studies[36–38].These data would require further investigation, but they provide an interesting first look at the molecular changes occurring during a repeated sonication regimen, which could provide additional benefits for the patients.

### Monthly repeated sonication shows promising results on pathological hallmarks

Finally, we assessed the effect of repeated sonication on amyloid deposits in the ARTE10 model (Fig4A). The ARTE10 mouse model is characterized by the presence of amyloid and mineralized deposits in the brain (Supp fig 4A), a noticeable loss of hippocampal neurons (Suppl Fig4B), and the presence of amyloid deposition around the brain vasculature (Suppl Fig3C). Consistently across all analyses, our data showed a significant reduction of the amyloid load in the very sparse sonication regimen (Group 6), consisting of a monthly sonication, compared to the other sonication regimens (Group 3 and 4, respectively p=0.0216 and 0.0437), with more regular sonication regimen. While not significant, it is worth noting that Group 6 also showed a slight reduction in the plaque area distribution compared to the non-sonicated control (Group 6 vs Group1, p=0.019, q=0.0385). Indeed, if analyzed separately, the average size of amyloid deposits was significantly reduced, while the percentage of amyloid positive area showed a tendency to decrease under the monthly sonication regimen (Supp Fig4D, p=0.1143, q=0.1154). The treatment did not have any effect on the overall number of deposits. Taken together, these data confirm the results previously shown that ultrasound alone can have beneficial effects in the treatment of Alzheimer’s pathology.

## Discussion

Altogether, this short pilot study showed that LIPU is a safe method to open the BBB, allowing repeated use with potential benefits in a pathological context. We previously demonstrated in a small cohort of nine patients with mild AD that seven repeated, bi-monthly LIPU-mediated BBB openings reduced the amyloid burden by 6.6% in the brain region targeted by the implantable ultrasound system used in the study[44]. The results from this preclinical study are in agreement and point to a modest effect of ultrasound alone on amyloid pathology. Further analyses of the different types of amyloid aggregates showed that repeated sonication triggers a reduction in the amyloid deposit size and amyloid distribution in the brain. These results are encouraging and point out the potential for ultrasound-mediated BBB opening as a therapeutic tool by itself. Further research should focus on finding the best synergistic effects between a specific therapeutic compound and an optimized sonication regimen to improve the outcome for patients.

Several studies have already explored the effect of sonication on amyloid pathology in AD models (see tables 4 and 5). However, the experimental designs between these different studies, including this one, make it complicated to grasp the real value of sonication in this pathology. In comparison to these models, ARTE10 model presents an amyloid deposition at 3 months, synaptic loss at 3 months, and a cognitive impairment at 12 months[47]. This model presents similarities with the TgCRND8 models. However, in the previously published studies, the animals were treated at a younger age (4.5mo in Jordao et al, 2010, 4mo in Jordao et al 2013, 6 to 8 mo in Burgess et al, 2014a and b, 7 mo in Poon et al, 2018). Our study explored the effect of repeated sonication in 1yo mice, where the amyloid deposition is already highly established and with a heavy load of amyloid deposits. 3TG and APP/PS1dE9 models were used at later stages, but the rate of amyloid deposition appears slower than in the ARTE10 mouse model (8mo in Shen et al, 2020; 2.5mo, 1yo and 2yo in Noel et al, 2023 and 14mo in Karakatsani et al, 2023, 12 and 14 mo in Hsu et al, 2018 and 16, 24 and 30 mo in Sun et al, 2021). To focus on comparable studies, we will only consider studies which used repeated sonication protocol to show the effect on the amyloid pathology[22,24,27–30,35]. Despite these experimental differences, similar effects were recorded, including our study: 1) Repeated sonication consistently triggers a decrease in the amyloid load across all models[22,24,27–30,35]; 2) A significant improvement of the cognitive performance[22,24,27,30], 3) An increased microglial activity around the deposits[22,29].

**Table 1:**
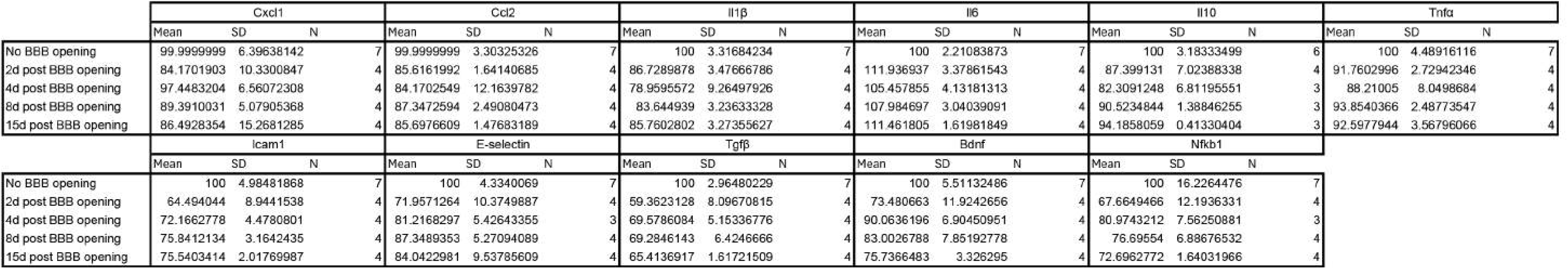
Row Statistics for the heatmap (Fig1D): Table summarizing for each group the mean, SD and N for the different markers analyzed. Triplicates were averaged per mouse.

**Table 2:**
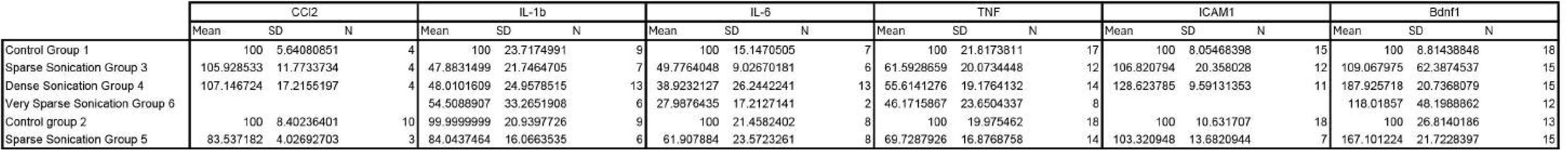
Row Statistics for the heatmap (Supp Fig3D): Table summarizing for each group the mean, SD and N for the different markers analyzed. As several data were missing, we analyzed Nplicates from different mice, but did not express the data per mouse for each group.

**Table 3:**
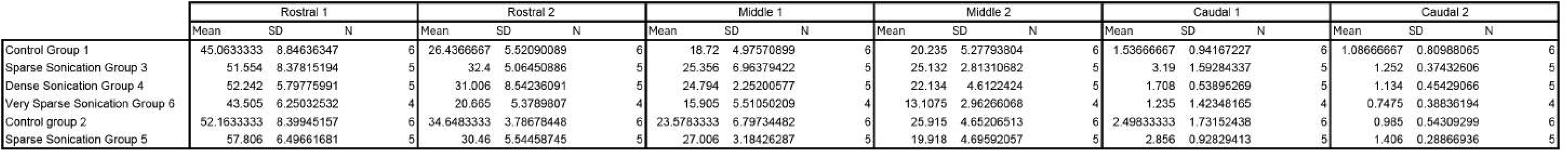
Row Statistics for the heatmap (Fig4C): Table summarizing for each group the mean, SD and N for the different locations analyzed. Triplicates were averaged per mouse.

**Table 4:** List of the previous mouse models used in combination with Ultrasound. Four models have been used to explore the effect of sonication on the amyloid pathology.

| Mouse model | Strain characteristics | Pathological hallmarks | Publications |
| --- | --- | --- | --- |
| 3TG model | Mix model for amyloid and tau pathologies<br>Single mutations in APP, MAPT, PSEN2 | • Cognitive impairment at 4 months,<br>• LTP deficits at 6 months,<br>• Amyloid deposition after 6 months,<br>• Tau pathology after 1 year | [22–24,83] |
| TgCRND8 model | Pure amyloid model<br>Double mutation in APP | • Amyloid deposition at 3 months<br>• Associated with cognitive decline<br>• Changes in LTP at 6 months | [4,25–28,84] |
| 5xFAD model | Pure amyloid model<br>Triple mutation in APP and double mutation in PSEN1 | • Early amyloid deposition around 2 months<br>• Synaptic loss after 4 months<br>• Associated with cognitive impairments<br>• Changes in LTP and neuronal loss at 6 months | [29,30,85] |
| APP/PS1dE9 model | Pure amyloid model<br>Single mutations in APP and PSEN1 | • Early LTP impairment at 3 months<br>• Synaptic loss at 4 months<br>• Amyloid deposition at 6 months<br>• Cognitive impairment around 1 year | [33,35,86] |

**Table 5:** Previous studies using ultrasound in mouse models for Alzheimer’s Disease and their output.

| Study | Model used | Age of the model | Sonication treatment | Associated therapeutics | Effect on the pathology | Effect on behavior |
| --- | --- | --- | --- | --- | --- | --- |
| Shen et al, 2020[22] | 3TG | 8 mo | Repeated, twice a week for six weeks | none | • Decreased amyloid burden in Cortex, CA and amygdala<br>• Reduced Tau phosphorylation<br>• Reduced branching and process length in microglia<br>• Axonal protection | Improvement in Y maze, Morris water maze and step down avoidance test |
| Noel et al, 2023[23] | 3TG | 2.5mo<br>1 yo<br>2 yo | One treatment | none | Opening volume slightly higher in 3TG compared to WT | NA |
| Karakatsani et al, 2023[24] | 3TG | 14 mo | Repeated, once a week for four weeks | none | >50% reduction in number and 42% in size | Improvement of memory (Morris water maze) |
| Jordao et al, 2013[25] | TgCRND8 | 4 mo | One treatment | none | • Reduced deposits in cortex<br>• IgG & IgM to penetrate infiltration<br>• Increased in Iba1 and GFAP<br>• Increased distal localization of microglia and astrocytes<br>• Increased neuronal area<br>• Increased colocalization between glial cells and amyloid deposits | NA |
| Burgess et al, 2014a[27] | TgCRND8 | 7 mo | Repeated, once a week for 3 weeks, bilateral | none | • Decreased in deposit size (~20%) and deposit number (~19%)<br>• Increased in DCX positive neurons (188% for nTG, 252% for Tg)<br>• Increased in dendrite length and branching | Improvement at the Y maze |
| Burgess et al, 2014b[26] | TgCRND8 | 6-8 mo | One treatment | none | • Correlation of vessel diameter and leakage probability<br>• Vessels displaying leakage are significantly smaller in Tg mice<br>• Vessels coated with amyloid show less<br>changes in diameter | NA |
| Poon et al, 2017[28] | TgCRND8 | 7 mo | Repeated, once a week for 5 weeks | none | Decreased number of amyloid deposits<br>Decrease amyloid deposit area | NA |
| Bobola et al, 2020[29] | 5xFAD | 6 mo | Repeated, daily for five days | none | Decreased amyloid burden by ~50% compared to control<br>Increased microglia colocalization with amyloid deposits compared to control | NA |
| Lee et al, 2020[30] | 5xFAD | 4 mo | Repeated, once a week for six weeks | none | Decrease of both amyloid deposits area and deposit area<br>Increased CSF abeta | Improvement in working memory |
| Jordao et al, 2010[4] | TgCRND8 | 4.5 mo | One treatment | BAM10 antibody | Slight decrease in Abeta deposits | NA |
| Alecou et al, 2017[31] | Rabbit, high cholesterol diet | Not disclosed | Repeated, three treatments | BC-10 (sigma), anti Amyloid-beta antibody | Original amyloid deposit density 200/cm<sup>2</sup>, decreased to 78/cm<sup>2</sup> after treatment | NA |
| Hsu et al, 2018[35] | APP/PS1dE9 | 12-14 mo | Repeated, once a week for five weeks | GSK-3 Inhibitor | Significant decrease of the deposit size in hippocampus, including with FUS alone | NA |
| Sun et al, 2021[33] | APP/PS1dE9 | 16 mo, 24-30 mo | Three treatments for 16mo cohort, one treatment for 24-30mo cohort | anti-pGlu3 (07/2a) | Decreased hippocampal plaque deposition in APP/PS1dE9 mice co-treated with anti-pGlu3 Aβ mAb and FUS-BBBD<br>Age-related hippocampal synaptic degeneration is reduced in APP/PS1dE9 mice after treatment with 07/2a and FUS-BBBD<br>Altered plaque-associated microglia and macrophage activation in aged APP/PS1dE9 after treatment with 07/2a and FUS-BBBD | No significant effects in fear learning and memory or locomotor activity with anti-pGlu3 Aβ mAb or FUS-BBBD |
| Deng et al, 2021[34] | APP/PS1dE9 | 10 mo | One treatment | IV injection of US-HA-Exo | Decreased deposit area and occupancy | NA |
| Antoniou et al, 2024[32] | 5xFAD | 5 mo | One treatment | Amyloid Protein (1–40) antibody produced in rabbit whole antiserum, A8326 | Allow for non-invasive antibody delivery to amyloid deposits | NA |

Our study demonstrated that in both a wildtype and a mouse model for AD with amyloid pathology (ARTE10), repeated LIPU-mediated BBB opening does not affect the general physiology of the mouse. However, we observed that a single sonication led to an increase of *Il-6*, while most of the other factors were either unchanged or slightly decreased (Fig.1D). *Il-6* is classically described as a pro-inflammatory cytokine with several key functions in the central nervous system but also shows anti-inflammatory functions in specific contexts[48]. *Il-6* has been shown to play a major role in activating microglia and favoring microvasculature repair after injury[49]. In brain diseases, *Il-6* expression was found to be increased in traumatic brain injury (TBI) as well as some initial steps of Alzheimer’s disease[50–52], however with conflicting associated functions. While the increase of *Il-6* levels after TBI was associated with anti-inflammatory and pro-survival effects, sustained elevated levels of *Il-6* were shown to be detrimental by increasing neuronal excitability leading to potential seizures[48]. Moreover, *Il-6* stimulates angiogenesis following a stroke episode, highlighting its potent role in stimulating vasculature repair after vascular injuries[53]. In addition, *Il-6* is known to increase in response to tissue injuries and to induce hematopoiesis (increase in red blood cell, Fig1B)[45]. In our experiments, we hypothesized that the increase in *Il-6* was tightly correlated to the vasculature repair process, without triggering a pathological chronic inflammatory response. Indeed, other cytokines and chemokines were unchanged or decreased after BBB opening, pointing out the unique role of *Il-6* in this context.

Other factors like the *Tnf-a*, another known partner involved in pathogenesis of AD[54], showed a downregulation post BBB-opening. Interestingly, in the context of TBI, *Il-6* inhibits the production of *Tnf-a* as an anti-inflammatory process, as observed in our data[48]. *Tnf-a* plays a central role in neuroinflammation[55], which reinforces the observation that LIPU-mediated BBB opening does not trigger pathological inflammation. Similarly to *Tnf-a*, *Il-1b* is associated with a myriad of negative effects on the brain, in regard to seizures, a role in the development of different psychiatric disorders and neurodegenerative diseases[56–58]. More importantly for the BBB, the inhibition of the *Il-1b* pathway showed a positive effect on the brain vascularization[59]. Counter intuitively, we observed a decrease in level of *Il-10*, an anti-inflammatory cytokine of the brain, with important roles in the repair process of the brain[60] and limiting pathological inflammation[61]. This suggests that LIPU-mediated BBB-opening is not a major traumatic event in the brain and only triggers transient changes in inflammatory markers in the brain. Our results showed a transitory effect on astrocytic activation (Fig. 3F). These results are in accordance with recent publications, showing a transient activation of astrocytes and microglia in the first hours post sonication, with a normalization to baseline after few days[46,62–64].

In this direction, the decrease in *nfkb* expression confirms the minimal impact of our BBB opening strategy on neuroinflammation[65,66]. Similarly, Ccl2, a factor known for disrupting the BBB integrity[67–69], as well as favorizing the infiltration of leukocytes and monocytes to the brain parenchyma[67,70], was found decreased post sonication, reinforcing the idea that LIPU-mediated BBB opening is a mild disruption of the BBB itself, and does not trigger pathways observed in stroke- or infection-mediated BBB disruptions. Our study also showed a transient decrease in surface adhesion molecule expression *Icam* and *E-Selectin*. *E-Selectin* levels were recovered 8d post opening, while *Icam* took longer to recover. In a recent publication, it was shown that repeated sonication schedules could be favorable for the vasculature, by stimulating higher level of adhesion molecule VE-Cadherin [71]. To really understand the effect of repeated sonication on the vasculature, especially in the context of neurodegenerative diseases, more in-depth studies will be required in the future, using high spatial resolution techniques such as in vivo 2-photon microscopy and spatial transcriptomics and proteomics. These studies could bring evidence of the beneficial effect of sonication in stimulating the repair process of the vasculature, and reperfusion of the brain[62].

Our study offers a unique timeline into the changes happening post BBB opening. Several studies attempted similar quantifications, however with a shorter timeline going up to 24h post sonication, versus 15 days in our study[72–74]. The data obtained 24h post sonication[72,73] showed a significant decrease in the magnitude of changes observed in the different factors, reaching similar amplitudes as what was observed in our study (0.5 to 1-fold change). Compared to these studies, our data showed that there are no negative long-term changes in induced neuroinflammation post sonication, confirming the normalizing level of all analyzed cytokines and chemokines in the previous studies. These results are a testimony to the transitory aspect of the US/MB-mediated BBB opening. It is however interesting to compare the data generated in our wildtype cohort, where a single BBB opening was performed, to the data obtained in our ARTE10 cohort, where different repeated sonication regimens were applied. Two main differences were observed between the two cohorts: The absence of *Il-6* increase and increase in *bdnf* expression in the repeated sonication cohort compared to the single sonication cohort. Increased *bdnf* was already observed post sonication[72] and suggested as the underlying mechanism responsible for stimulating neurogenesis in other studies[27,36–38]. For example, in Kovacs et al., *bdnf* levels were observed to increase around 24h post sonication. Our study showed that this increase might be transient, as we did not observe a detectable increase at 48h post-sonication or could be an effect of the much stronger sonication parameters used to disrupt the BBB in the Kovacs et al. study. However, as we observed an increase of *bdnf* levels after repeated sonication, it suggests that regular opening of the BBB could allow sustained levels of BDNF overtime, which could in turn have a beneficial effect, potentially through increased neurogenesis or higher neuronal excitability and synaptic homeostasis[75]. However, as observed for groups 3 and 4, higher levels of BDNF did not correlated with decrease of amyloid pathology. It is now well documented that neuronal activity correlates with amyloid beta secretion in the intercellular spaces[76–78]. These results would suggest that LIPU-mediated BBB opening might efficiently regulate amyloid beta pathology within a certain range of stimulation.

Cerebral amyloid angiopathy (CAA), characterized by the amyloid deposition around the blood vessel, is found in most patients presenting Alzheimer’s disease. The accumulation at the vessel is believed to originate from incomplete clearance of the amyloid from the brain. Previous work has established that accumulation of amyloid in the blood vessels progressively replaces all the different tissues forming the arterial wall[79]. Accumulation of amyloid in the arterial wall affects blood vessel integrity and function, resulting in cerebral infarct or hemorrhage[80]. Post sonication, it was shown that vessels containing amyloid were less susceptible to diameter changes during US/MB-mediated BBB opening protocol, suggesting a more rigid structure. In addition, in a mouse model for AD, small vessels were most likely to display leakage post BBB opening in their littermate controls[26]. Unfortunately, the presence of CAA in patients with AD could be the reason for ineffective anti-amyloid immunotherapy and observed adverse events leading to microhemorrhage in patients[81,82]. In our study, the ARTE10 model also presents some level of CAA, with notable accumulation of amyloid positive staining around the blood vessels (Supp Fig. 3C, Supp fig.4). Targeted analysis did not show any beneficial effect of repeated sonication on the vascular amyloid (Fig 4). However, it is important to note that postmortem brain imaging did not reveal the presence of increased hemorrhage following sonication. This suggests that sonication can still be a safe and potent strategy to efficiently deliver therapies against amyloid to the brain [23]. As mentioned earlier, few studies have already explored this possibility[4,31–33,35], including during a clinical trial[15]. All studies reported a beneficial effect of therapeutic delivery to the brain and a reduction of the pathology in the different models used (See tables 1 and 2).

**Figure 4:**
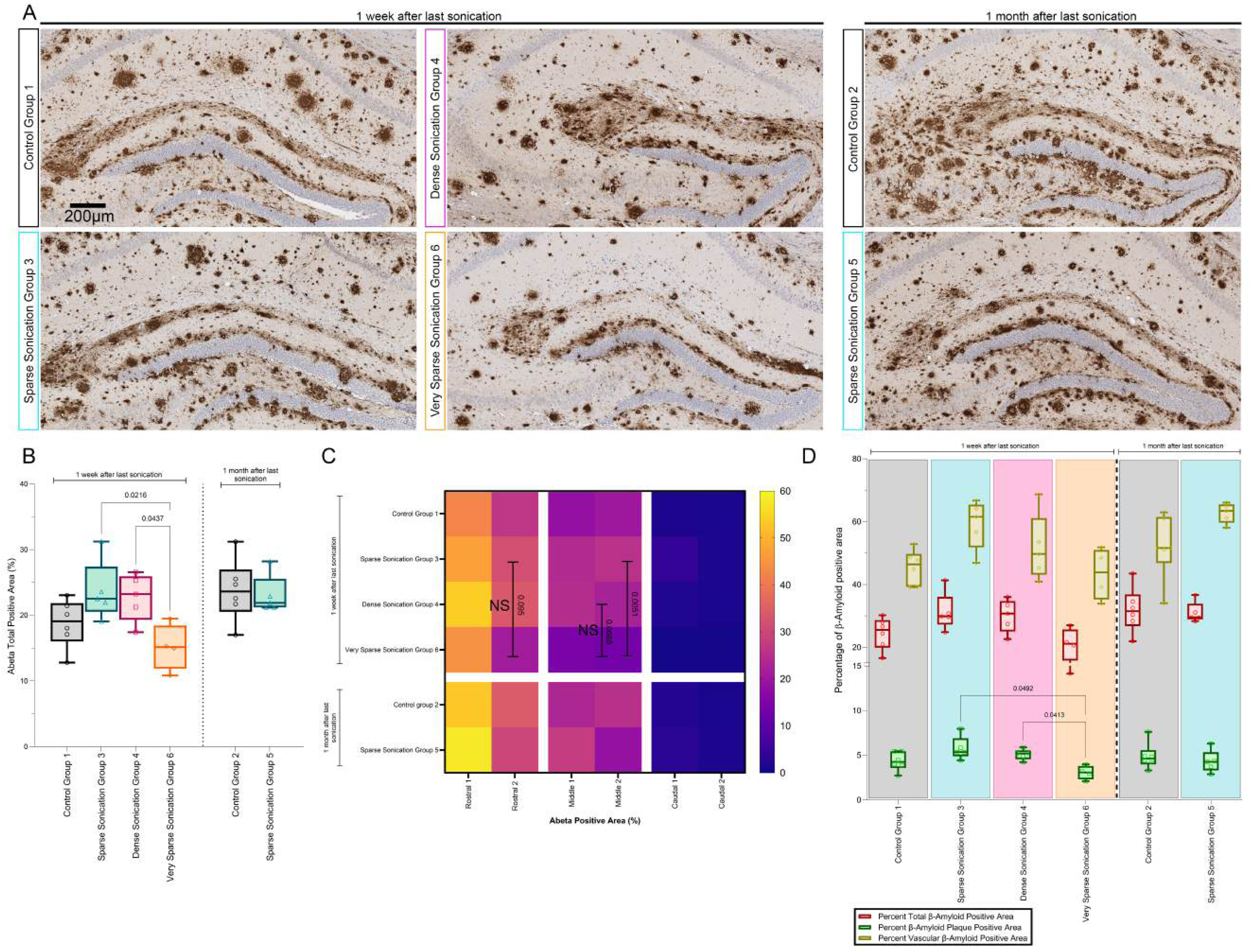
Decrease of the amyloid load following monthly sonication regimen: **A)** Representative images of the hippocampal region isolated from immunohistochemistry staining of b-amyloid on coronal sections. **B)** Box plot showing median, 25 and 75 percentiles and min/max values, along with individual data point for each animal representing the total amyloid load (in percentage of positive area) for each group. Each dot represents an independent animal from the group. One-way ANOVA with post hoc Tukey’s multiple comparisons test, F(3, 16)=0.04014, p=0.0163. **C)** Heatmap representing the level of amyloid deposition in the brain at different locations from rostral to caudal. Pseudo coloring represents the Median of each group. 2-way ANOVA, difference between groups F (5, 25) = 3.323, P=0.0195. D) Box plot showing median, 25 and 75 percentiles and min/max values, along with individual data point for each animal, for the total amyloid positive area (red boxes), amyloid deposits (green boxes) and vascular amyloid deposition (yellow boxes). 2-way ANOVA with post hoc Tukey’s multiple comparisons test, difference between groups F (5, 25) = 4.317, P=0.0057.

## Conclusion

Altogether, this study brings additional evidence to the field that repeated sonication to disrupt the BBB is a safe procedure, which could bring positive therapeutic effects in the context of neurodegenerative diseases by both enhancing drug delivery and stimulating mild, transient inflammatory responses. More in-depth analyses are required to understand the molecular and cellular mechanisms at play. However, it is worth considering the potential therapeutic effect of ultrasound mediated BBB opening in the future, and to explore the synergic effects of their combination to disease modifying therapeutics, which could lead to improved outcomes for patients.

## Acknowledgements

The authors acknowledge Inotiv (former Bolder BioPATH) for their work on this study as well as Dallas Tissue Research for their help with image analysis. We want to thank Prof. Benoit Delatour for his input on the paper. This study was fully funded by Carthera (Lyon, France).

## Contributions

M.C. and A.C. designed and supervised the study. M.C. and G.B. developed the preclinical set-up for BBB opening, G.B. provided technical expertise. P.D.R. performed data curation, formal analyses and statistical analyses. M.C. and P.D.R prepared the original manuscript and the revised manuscript. All authors attest that no generative AI or AI-assisted technologies were used in the writing process.

## Data Availability statement

Data presented in this paper are available on request to the corresponding author.

---

**Figure S1:**
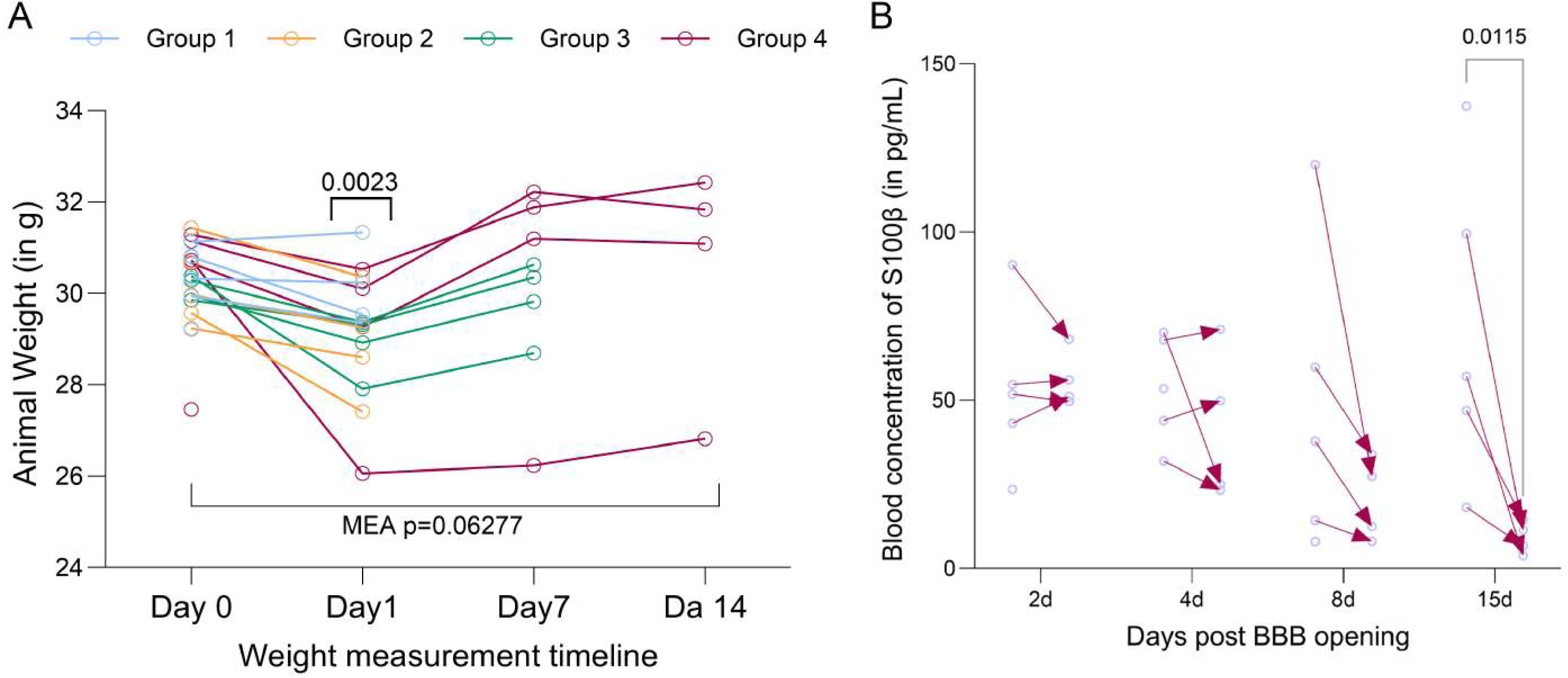
Health check and BBB opening markers: **A)** line graph representing the evolution of the mouse weight for each group. Each line represents an individual mouse. Group 1 (blue) and group 2 (orange) only have 2 measurements, with a respective sacrifice at day 2 and day 4 post sonication. Mixed effect analysis with post hoc Sidak’s multiple comparisons test, difference between timepoints F (0.1883, 1.569) = 5.678, P=0.06277, D0 vs D1 p=0.0023, **B)** Arrow plot representing the blood level of S100β in samples 15min post sonication and at time of sacrifice (24h, 48h, 8d and 15d post sonication). Direction of the arrow indicates the evolution of the level (ie downward arrow shows decreasing level, upward arrow shows increasing level). Mixed effect analysis with post hoc Sidak’s multiple comparisons test, difference between timepoints F (1, 12) = 10.91, P=0.0063, post sonication vs terminal 15 days p= 0.0115.

**Supplemental Figure 2:**
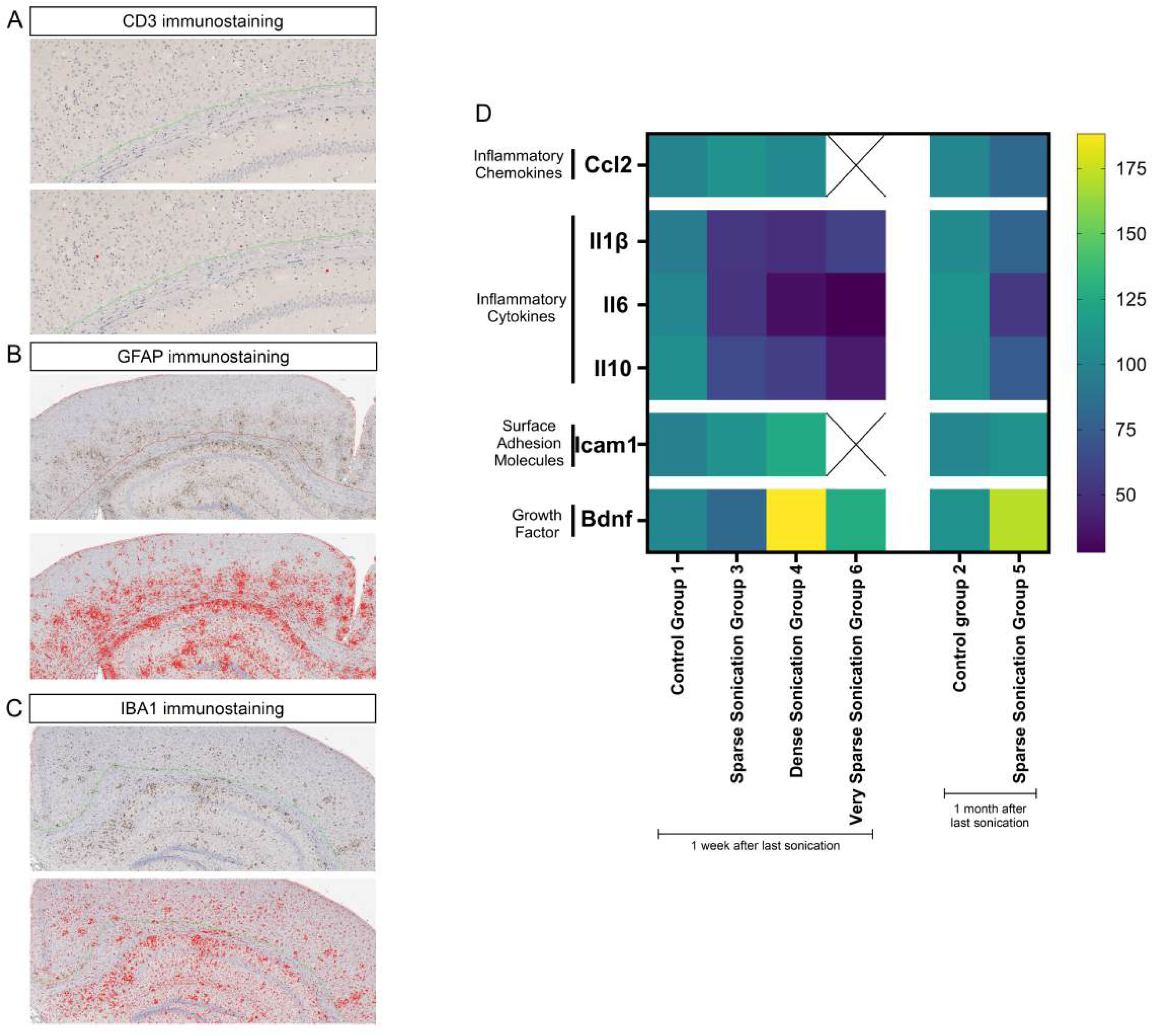
automated image analysis pipeline and inflammatory marker expression post repeated sonication protocol: **A-C)** Representative example of the automated pipeline allowing detection of CD3+ cell **(A)**, GFAP **(B)** and Iba1 **(C)**. Positive cells are highlighted in red, allowing the quantification of the number of CD3+ per mm2 and the surface of occupancy for GFAP and Iba1 positive cells. **D)** heatmap representing the pilot RT-Q-PCR dataset collected at each timepoint and exploring the level of inflammatory markers post repeated sonication. No statistical analyses were performed due to the lack of independent data available.

**Supplemental Figure 3:**
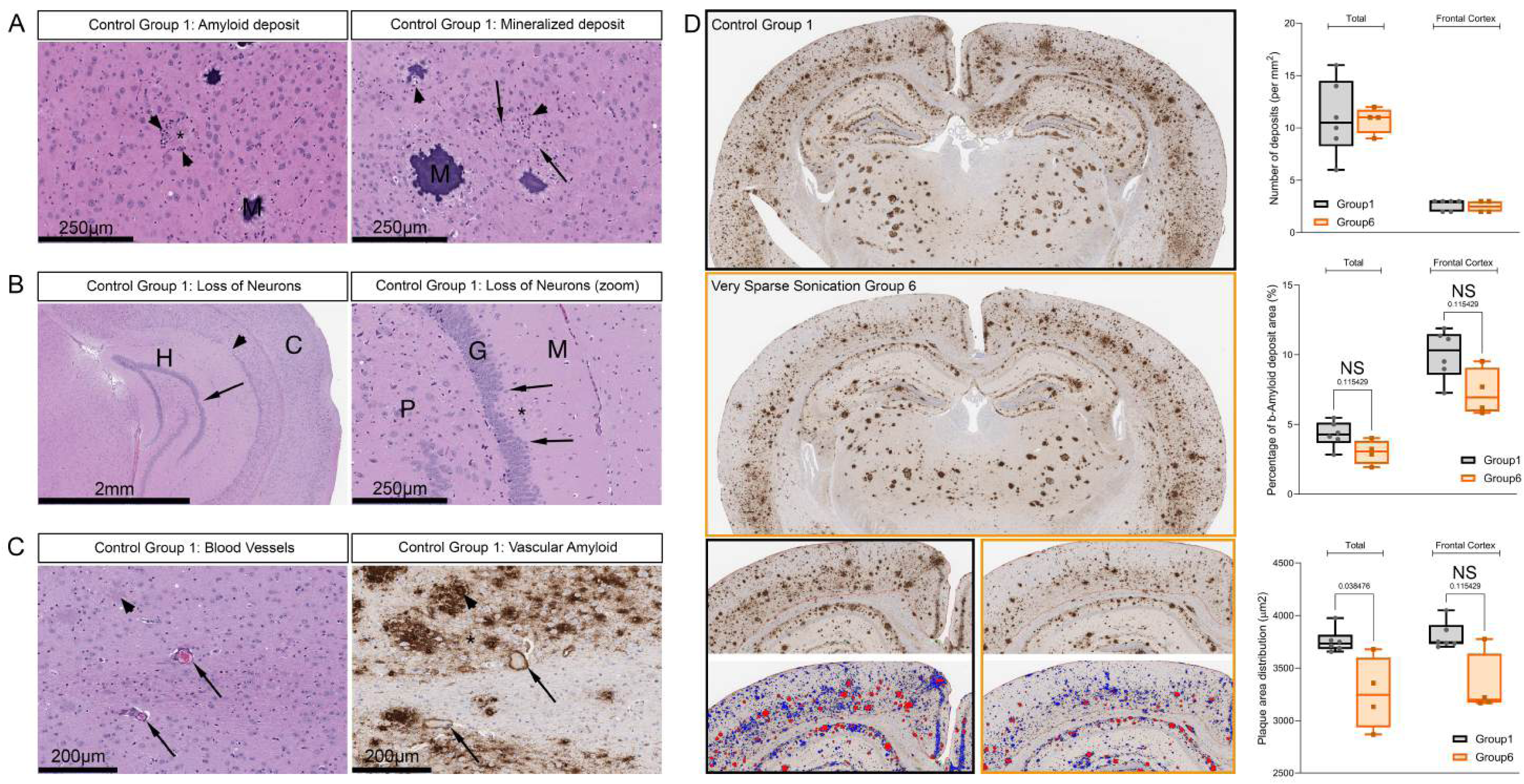
Pathology of the ARTE10 and supplemental analyses for the effect of a monthly sonication regimen: **A)** Representative images showing different types of pathological deposition, with the amyloid deposits (left) and the mineralized deposits (right, M). **B)** Representative images showing loss of hippocampal neurons in the cellular layer of the dentate gyrus. Arrows point at the area showing loss of neuronal cell bodies. H=hippocampus, C=cortex, M=molecular layer, G=granular layer, P=polymorphic layer. **C)** Representative images highlighting the presence of amyloid deposition around the brain vasculature. Arrows point at the vascular deposition, observed on serial sections. **D)** Side by side comparison of group 1 (control) and group 6 (very sparse sonication), highlighting the effect of the monthly sonication. Number of amyloid deposits (per mm2), percentage of amyloid positive area and amyloid deposits area was compared between the two groups. For all the comparisons, Group1 and Group6 were compared using nonparametric multiple t-test Mann-Whitney. Graphical representations show median, 25 and 75 percentiles and min/max values, along with individual data point for each animal, and statistical values showing calculated q-value. Percentage of b-amyloid deposit area, total, group 1 vs group 6, p=0.1143, q=0.1154; Frontal cortex, group 1 vs group 6, p=0.1143, q=0.1154. Average deposits area, total, group 1 vs group 6, p=0.019, q=0.0385; Frontal cortex, group 1 vs group 6, p=0.1143, q=0.1154.

**Supplemental Figure 4:**
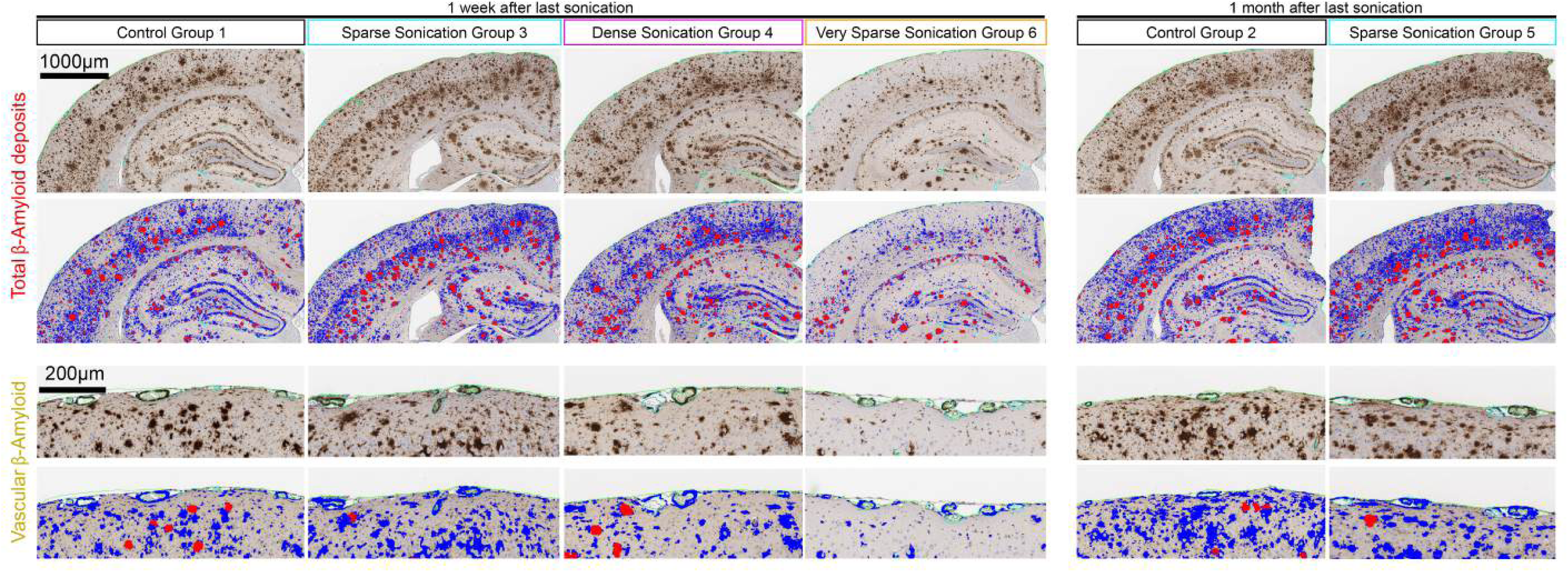
Artificial intelligence quantification of amyloid pathology: Representative images showing the total amyloid load by immunohistochemistry and pseudo color from the automated quantification pipeline, as well as the vascular amyloid deposition by immunohistochemistry and pseudo color from the automated quantification pipeline.

**Supplemental Figure 5:**
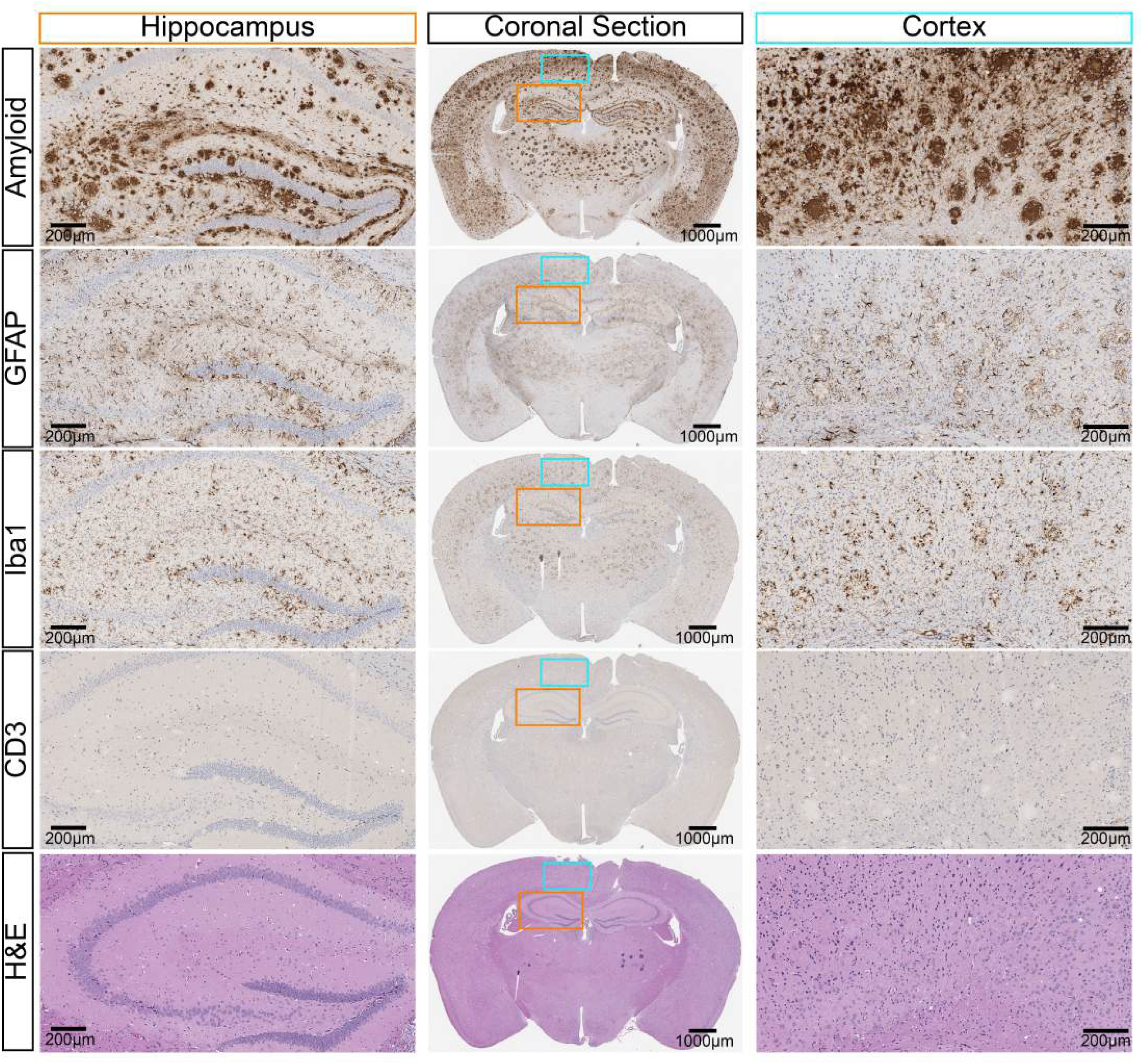
Full brain scans of each groups, respectively Group 1 – SuppFig5, Group 2 – Supp Fig 6, Group 3 – Suppl Fig 7, Group 4 – Suppl Fig 8, Group 5 – Suppl Fig 9, Group 6 – Suppl Fig 10

**Supplemental Figure 6:**
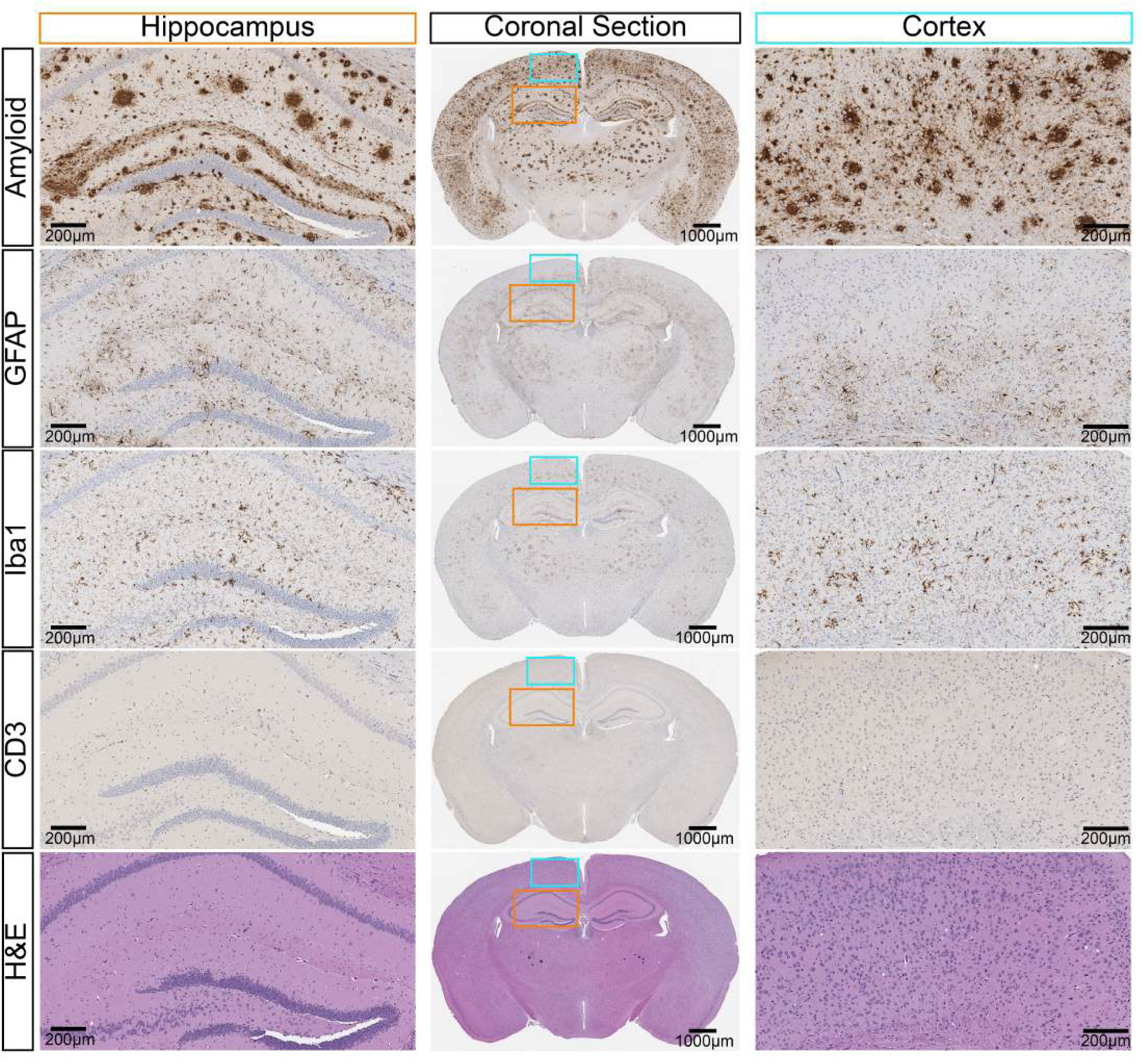
Full brain scans of each groups, respectively Group 1 – SuppFig5, Group 2 – Supp Fig 6, Group 3 – Suppl Fig 7, Group 4 – Suppl Fig 8, Group 5 – Suppl Fig 9, Group 6 – Suppl Fig 10

**Supplemental Figure 7:**
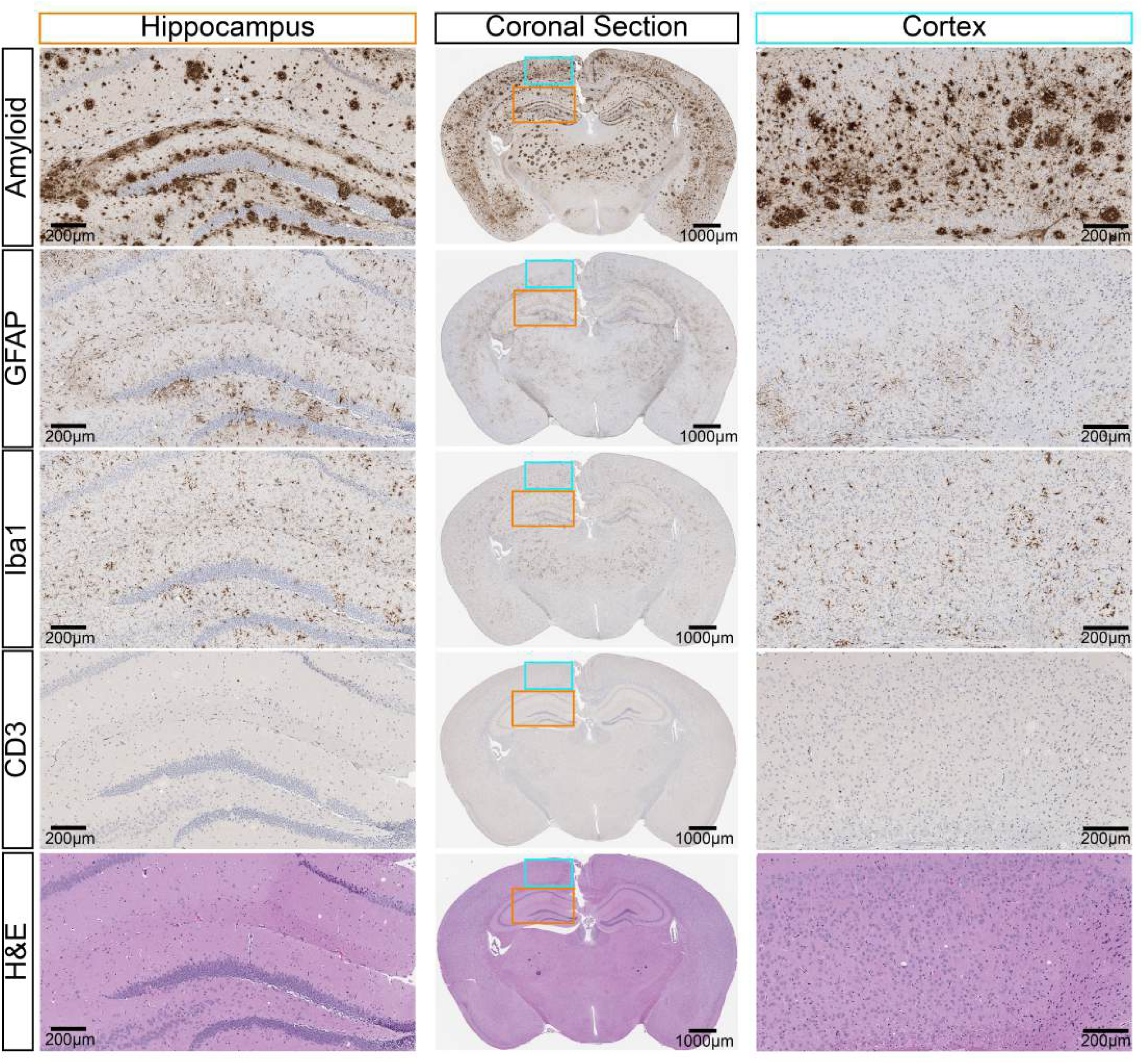
Full brain scans of each groups, respectively Group 1 – SuppFig5, Group 2 – Supp Fig 6, Group 3 – Suppl Fig 7, Group 4 – Suppl Fig 8, Group 5 – Suppl Fig 9, Group 6 – Suppl Fig 10

**Supplemental Figure 8:**
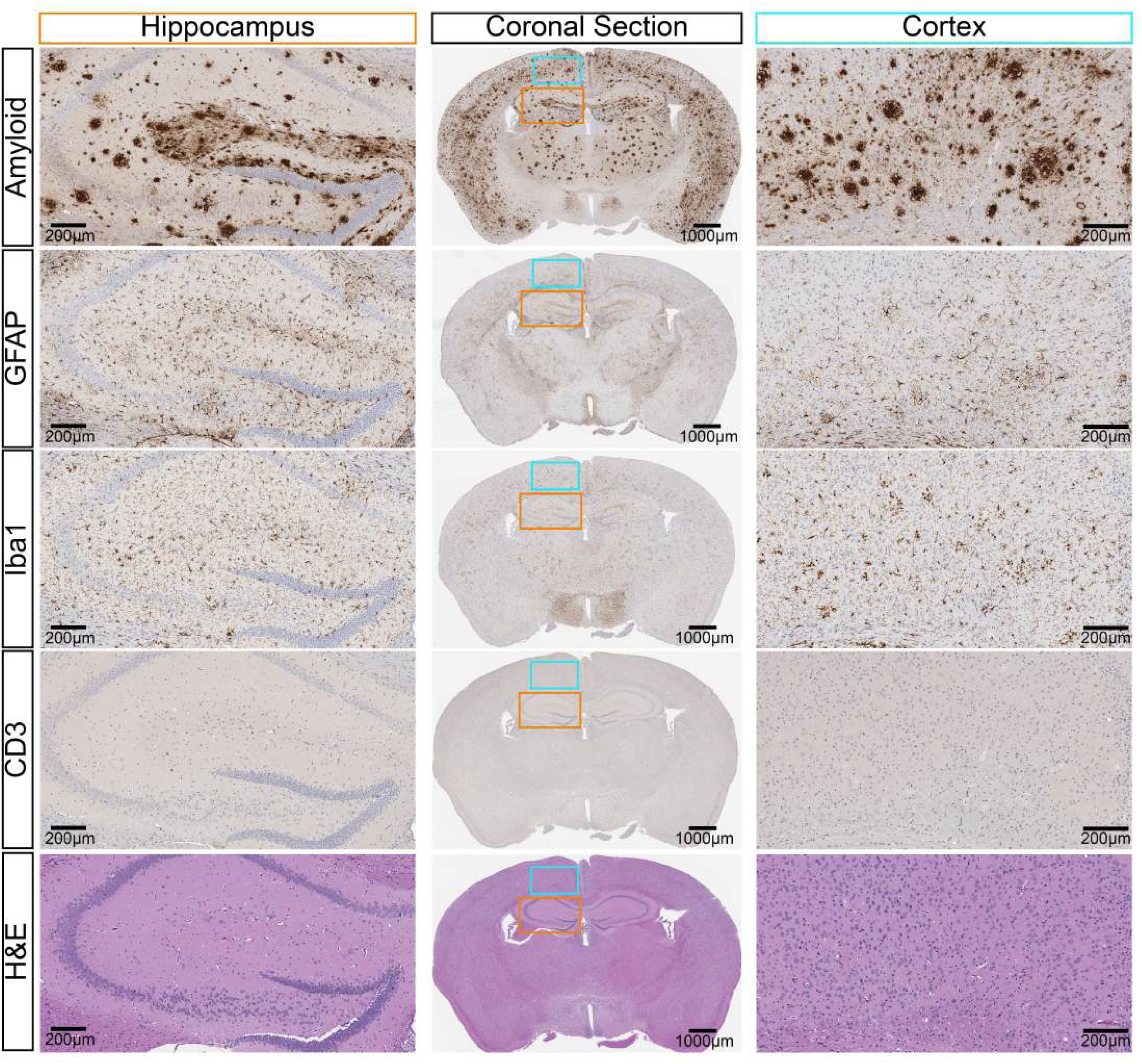
Full brain scans of each groups, respectively Group 1 – SuppFig5, Group 2 – Supp Fig 6, Group 3 – Suppl Fig 7, Group 4 – Suppl Fig 8, Group 5 – Suppl Fig 9, Group 6 – Suppl Fig 10

**Supplemental Figure 9:**
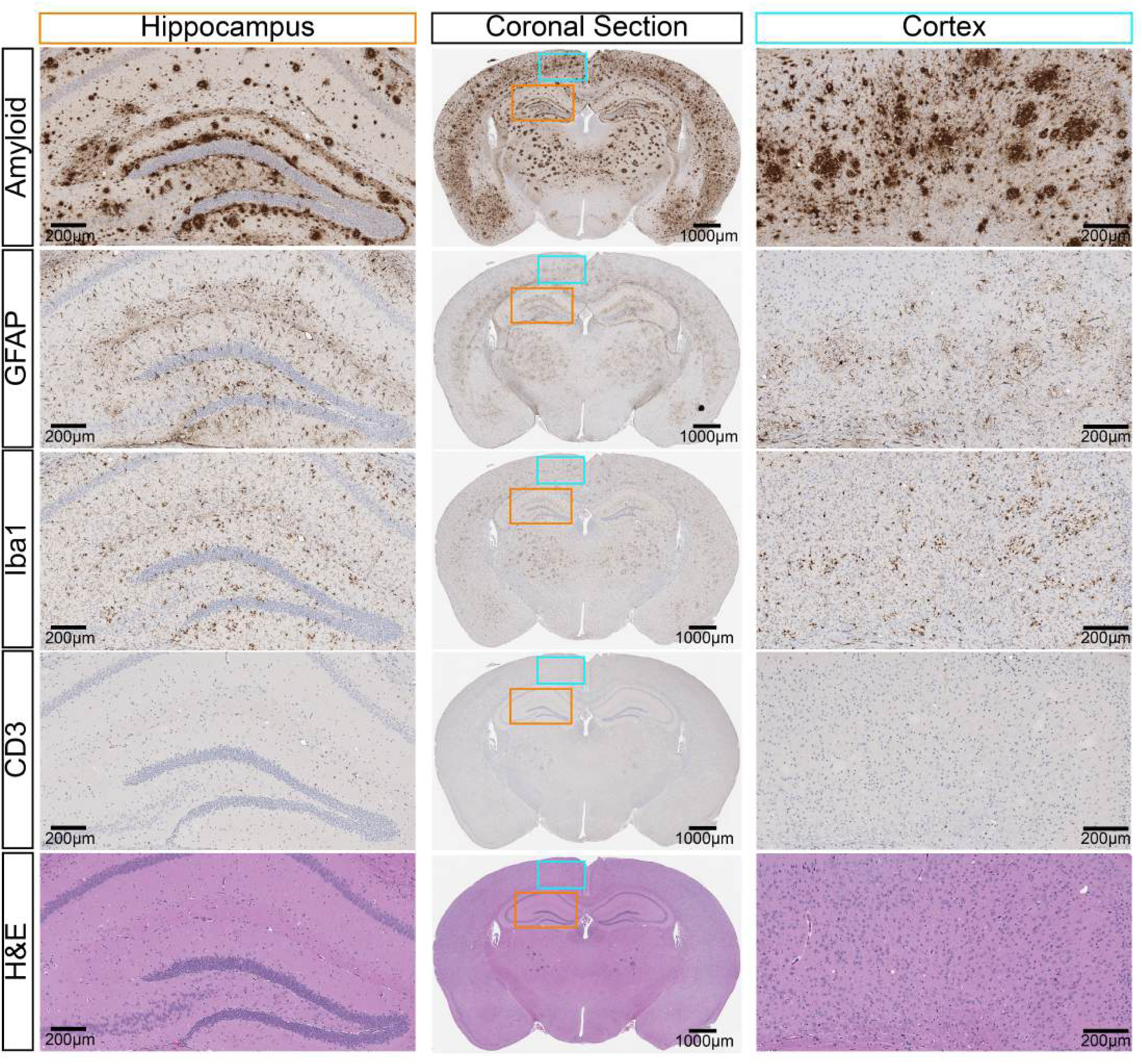
Full brain scans of each groups, respectively Group 1 – SuppFig5, Group 2 – Supp Fig 6, Group 3 – Suppl Fig 7, Group 4 – Suppl Fig 8, Group 5 – Suppl Fig 9, Group 6 – Suppl Fig 10

**Supplemental Figure 10:**
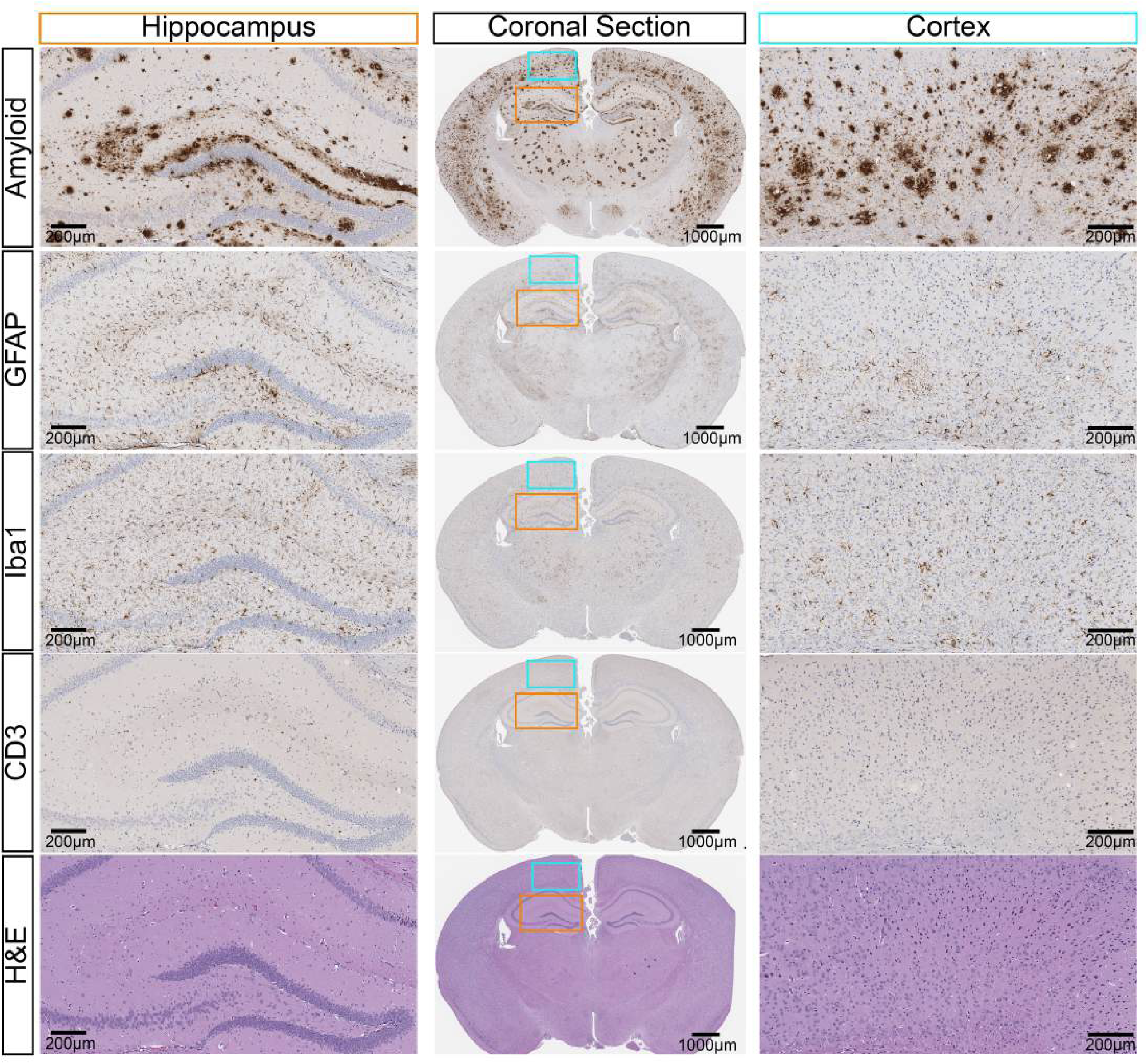
Full brain scans of each groups, respectively Group 1 – SuppFig5, Group 2 – Supp Fig 6, Group 3 – Suppl Fig 7, Group 4 – Suppl Fig 8, Group 5 – Suppl Fig 9, Group 6 – Suppl Fig 10

